# Evaluation of resistant starch from teff (*Eragrostis tef*) grain as a film coating material for colon-targeted drug delivery

**DOI:** 10.1101/2024.08.06.606940

**Authors:** Yohannes Teshome, Anteneh Belete, Tsige Gebre-Mariam

## Abstract

Teff (Eragrostis tef, family Poaceae) is a native cereal crop widely grown in Ethiopia, containing approximately 73% carbohydrates, of which about 30% is resistant starch. This study evaluates resistant starch extracted from teff grain as a film coating material for colon-targeted delivery of metronidazole, used as a model drug. Starch was extracted from teff and resistant starch was isolated from the total starch. Metronidazole core tablets were prepared by wet granulation, compressed, and coated with a resistant starch-based film. The physicochemical properties of the tablets were evaluated *in vitro*. To prevent premature film disruption caused by the swelling of amylose, a dominant component of resistant starch, a water-insoluble polymer, ethylcellulose, was added. Various proportions of amylose and ethylcellulose were used as film coating materials and evaluated in simulated conditions to determine the optimal combination for drug release in the colon, but not in the upper gastrointestinal tract. The results of the dissolution and fermentation studies indicated the best film coating proportions of amylose to ethylcellulose and the corresponding thicknesses in percentage of total weight gain were: 1:1 ratio at 6% thickness, 1:2 ratio at 4% and 6% thickness, and 1:3 ratio at 2% and 4% thickness. The targeted drug release of the film material is attributed to bacterial enzyme digestion of the resistant starch component in the colon. The digestion of resistant starch creates pores in the ethylcellulose film scaffold, leading to the disruption of the film and release of the drug exclusively in the colon, where the bacterial microflora reside. Based on these results, resistant starch from teff grain shows potential as a colon-targeting excipient.

## Introduction

Starch is a versatile excipient utilized in various pharmaceutical dosage forms. Traditionally known for its role as a disintegrant or diluent, starch is increasingly being explored as an advanced drug carrier. The therapeutic efficacy of starch-adsorbed, starch-encapsulated, or starch-conjugated drugs largely depends on the type of starch used. Its amenability to undergo physico-chemical modifications makes starch suitable for developing novel drug delivery systems, including targeted drug delivery systems [1, 2].

Despite numerous studies highlighting the expanded use of starch as an excipient in the pharmaceutical industry, there have been limited efforts to evaluate resistant starch (RS) for colon drug delivery systems. Targeting drug delivery to the colon offers several advantages, including the local treatment of various bowel diseases and systemic delivery of protein and peptide drugs. This approach increases efficacy and reduces the side effects of drug treatments [3]. Colon-targeting drug delivery systems (CTDDS) protect orally administered drugs until they reach the colon, preventing drug release and absorption in the stomach or small intestine. Various approaches for CTDDS include transit-time-dependent, pH-dependent, bacterial enzyme-dependent, and osmotic pressure-controlled systems. The bacterial enzyme-dependent CTDDS can be a prodrug-based, coating-based, or matrix-based system [4].

Chen *et al.* [1] developed a novel protein drug matrix tablet for colon targeting using RS as a carrier prepared by pre-gelatinization and cross-linking of starch. An oral colon-targeting controlled release system using RS acetate as a film-coating material has also been developed [5]. Meneguin *et al.*[6] used RS-pectin films intended for colonic drug delivery by blending retrograded starch with pectin as a dispersion film-forming material. Other studies have also reported the use of starch-based drug delivery systems for colon targeting [5, 7, 8].

Most native starches have limited direct applications due to their instability with respect to changes in temperature, pH, and shear forces. Therefore, native starches are often modified to develop specific properties such as solubility, texture, adhesion, and tolerance to heating temperatures in industrial processes [9].

Starch can be categorized as digestible (non-RS) or resistant starch (RS) based on its digestibility. Digestible starches are broken down by body enzymes and include rapidly digestible starches (RDS) and slowly digestible starches (SDS). In contrast, RS consists of dietary starch and starch degradation products that escape digestion in the small intestine of healthy individuals [10, 11]. Starches are made up of two polymers of D-glucose: amylopectin, which has highly b ranched chains with a larger surface area, and amylose, which is unbranched with a smaller surface area. The branched structure of amylopectin makes it more accessible to digestive enzymes, while the tight structure of amylose resists enzymatic digestion. The amylose content is responsible for starch’s film-forming, gelling, and binding properties [12].

Although starch films can be used in different drug delivery systems, amylose’s tendency to swell in aqueous media can lead to accelerated drug release in the upper gastrointestinal tract (GIT) before reaching the distal GIT. To control this swelling, commercially available controlled-release polymers such as chitosan, alginate, or ethylcellulose (EC) are mixed with amylose to prevent drug release in the stomach and small intestine [3].

In this study, hydrophilic amylose was mixed with hydrophobic EC in various proportions to evaluate its potential as a film-coating material for CTDD. The use of RS in CTDDS takes advantage of its digestion by colonic bacterial enzymes, rather than by enzymes in the upper GIT. This study focuses on a bacterial enzyme-dependent CTDDS using RS as a coating material.

Teff (*Eragrostis teff*), a cereal crop native to Ethiopia, has an attractive nutritional profile, comprising 13% protein, 2.5% total fat, 9% vitamins, minerals, and other components, and 73% carbohydrates. Approximately 20-40% of the total starch in teff is RS [13]. Unlike many other cereals, teff is gluten-free, making it suitable for individuals with gluten-triggered allergies or gluten-sensitive enteropathy, such as those with celiac disease [14, 15]. Therefore, using teff’s amylose in dosage forms can avoid allergic reactions in this population. The purpose of this study is to evaluate RS obtained from teff grains as a film-coating material for CTDD of metronidazole, used as a model drug.

## Materials and Methods

### Materials

#### Plant materials

Four varieties of teff grains, two “white” varieties (Debre-Zeit /DZ-01-196 and DZ-CR-37) and two “brown” varieties (DZ-01-1681 and DZ-01-99). These grains were obtained from the Holeta Agricultural Research Institute in Ethiopia. Additionally, a teff variety between white and brown, locally called "*sergegna*," was sourced from a local market in Addis Ababa for preliminary studies and larger-scale production.

#### Chemicals, enzymes and solvents

Pancreatic α-amylase (SOLARAY® Dietary Supplements, Park City, UT 84098 USA), Amyloglucosidase (AMG ASIN, LD Carlson Co. Ohio, USA), Pepsin (BCBR 3132v, Sigma-Aldrich/Switzerland), Pancreatin (Caelo Ch. Caesar & Loretz GmbH Pharma. Hilden, Germany), Sodium metabisulphite (BDH Chemicals Ltd Poole, England), Absolute ethanol (Carlo Erba Reagents, Italy), Metronidazole (Hubei Hongyuan Pharmaceutical Technology Co, Ltd. China), Ethylcellulose (BDH Chemicals Ltd Poole, England), Povidone K 29-32 (China Associate Group co., Ltd., China), Sodium starch glycolate (Huzhou Zhanwang Pharmaceutical Co. Ltd, China), Lactose monohydrate (Shreeji Pharma International, India), Propylene glycol (Horst G.F. von Valtier GmbH and Co. KG, Germany), Ethanol 96%, iodine solution and Erythrosine/Red No.3 (Kronos International Inc., Germany) were used as received.

## Methods

### Isolation of TS from teff

Teff grains were first sieved to remove extraneous materials, then milled using a laboratory mill (FRITSCH Pulverisette 2, RoHS, Idar-Oberstein, Germany) and sieved through a 315 µm sieve to produce whole grain meal. Small-scale extraction of TS followed the method described by Bultosa *et al.* [15], while larger-scale extraction was based on the method by Gebre-Mariam and Schmidt [16].

### Isolation of RS from the TS

The RS was isolated using the AOAC official method [17]. Briefly, 100 g of TS was suspended in 250 ml of distilled water to form a slurry. Pancreatic α-amylase and amyloglucosidase (AMG) were added, and the mixture was stirred for 16 h at 37°C to solubilize and hydrolyze the non-RS to D-glucose. The reaction was terminated by adding 96% ethanol, and the RS was recovered as a pellet by centrifugation (8000 rpm for 10 minutes). The pellet was washed with 96% ethanol and centrifuged again. This enzymatic digestion process was repeated three times to ensure complete digestion of the digestible starch.

### Test for starch

Four test tubes labelled D1, D2, D3, and D4 were prepared with a few grams of specific starch powder from the four different teff varieties in distilled water. A fifth tube with distilled water only was used as a blank. A drop of iodine solution (0.2 g iodine and 2 g potassium iodide in 100 ml of distilled water) was added to each test tube, and the resulting colour was observed.

### Test for RS

Two methods exist for testing RS: a direct method that quantifies RS in the residues after removing digestible starch using the AOAC Official Method (15) and an indirect method that determines RS as the difference between TS and digestible starch [17]. In this study, the indirect method was used. After digesting the digestible starch with enzymes, the remaining pellets were considered RS.

### Preparing film coated colon-targeted metronidazole tablets using teff RS

Metronidazole core tablets were prepared by wet granulation method. The formula for these tablets is shown in Table 1.

**Table 1:**
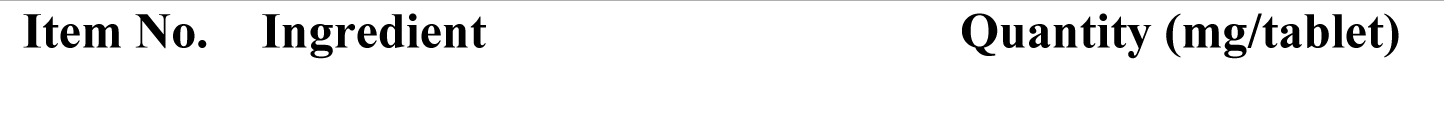
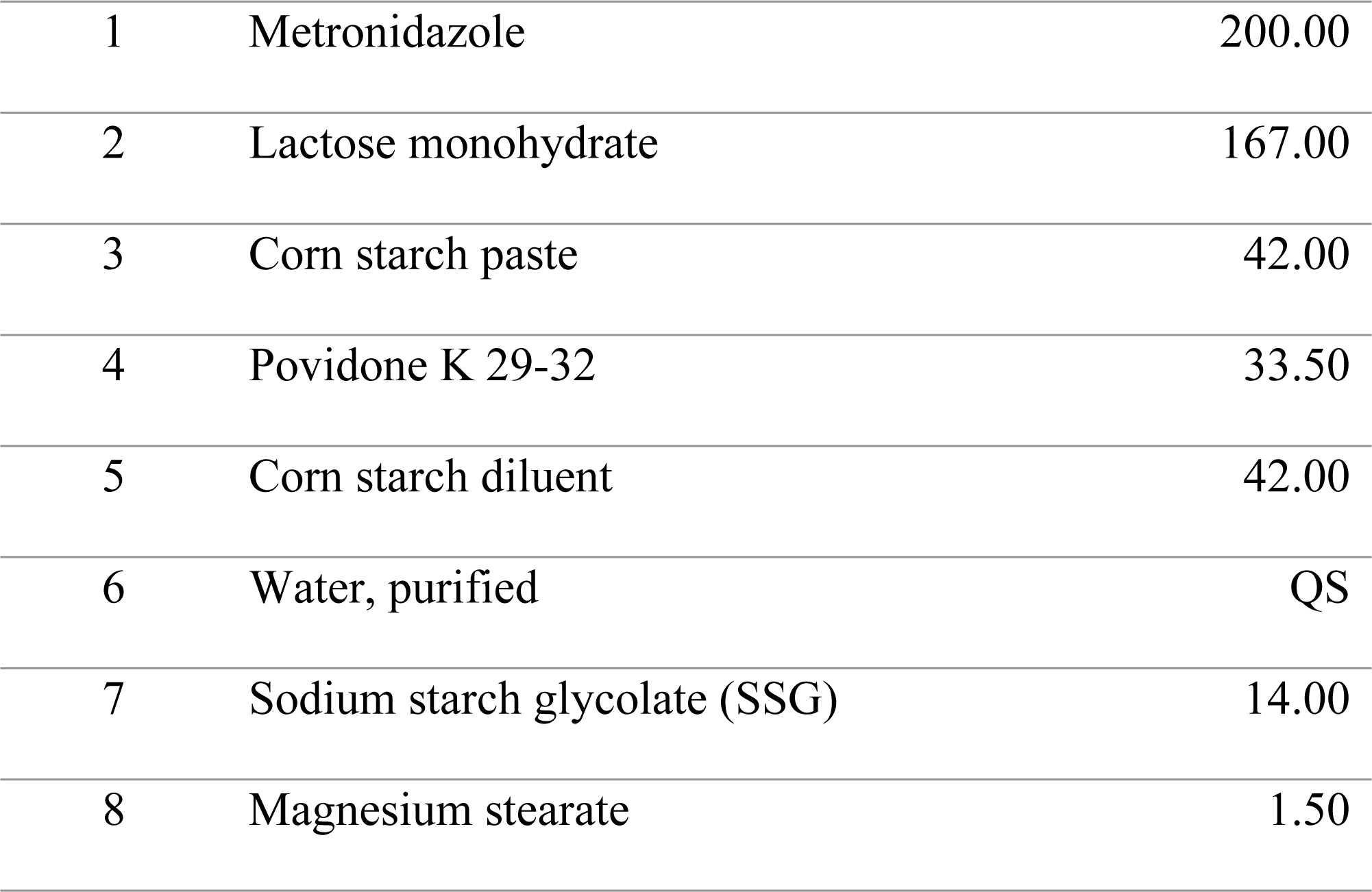
Metronidazole tablets (200 mg), formula for 1000 tablets.

### Preparation of corn paste

Corn starch paste (item 3, adhesive binder) was prepared by mixing corn starch with purified water (item 6, solvent) in a stainless-steel container. This mixture was heated over a boiling water bath and stirred until a translucent paste formed.

### Preparation of metronidazole core tablets

#### Ingredient processing

Metronidazole, lactose monohydrate (item 2, diluent), and corn starch (item 5, diluent) were passed through a 500-µm aperture screen. The ingredients were then transferred to a mixer (Machines Collette, Type: MP20, Model No: 87M2279; Keerbaan 70, Wommelgem, Antwerp, Belgium). Povidone was added to the mixer and mixed for 5 minutes.

#### Granulation

The prepared corn starch paste was added to the powder mixture and mixed until a suitable consistency was achieved. Additional water was added as needed. The wet mass was passed through a 2.5 mm sieve to form wet granules. The wet granules were spread on paper-lined trays and dried in an oven (KOTTERMAN®, Germany) at 50°C overnight. The dried granules were passed through a 1.5 mm aperture sieve and stored in an airtight container.

#### Compression of core tablets

The dried granules were transferred to a planetary mixer (Machines Collette, Type: MP20, Model No: 87M2279; Keerbaan 70, Wommelgem, Antwerp, Belgium). SSG (super-disintegrant) and magnesium stearate (lubricant) were screened through a 600-µm aperture screen and added to the blender. The mixture was blended for 5 minutes. The granules were compressed using a 10-station rotary tablet press machine (Rimek Mini Press-II, India) with 14/32-inch, standard concave punches at a pressure of 6.5 to 10.5 kg/cm². Each batch contained 200 mg of metronidazole with an average tablet weight of 500 mg.

### Evaluation of granules and core-tablets

#### Evaluation of the granules

Parameters such as angle of repose, flow rate, bulk density, and tapped density were measured using 30 g samples. Carr’s index and Hausner ratio were calculated.

#### Evaluation of the core-tablets

Physicochemical parameters such as weight variation, friability, hardness, diameter, thickness, and disintegration time were evaluated according to USP 30 [18].

### Calibration curve

A series of metronidazole solutions were prepared in appropriate media, and their absorbance was determined using UV spectrophotometry at a λmax of 278 nm. Calibration curves (concentration vs absorbance) were plotted in simulated gastric (pH 1, 0.1 N HCl, 0.32% pepsin), small intestinal (phosphate buffer, pH 6.8, 1% pancreatin), and colonic environments (phosphate buffer, pH 7.2, with 10% human faeces).

### Optimization of film forming property of amylose (RS) with ethylcellulose (EC)

#### Preparation of film coating solution

Various ratios of amylose: EC (1:1, 1:2, 1:3, 1:4, 1:5) were prepared as described elsewhere [19, 20]. Propylene glycol (plasticizer) and erythrosine (colorant) were added at 2.5% and 0.3%, respectively. A control film coating solution with only amylose and plasticizer was also prepared (1:0 ratio).

#### Amylose solution preparation

10 g of amylose was dissolved in 100 ml of 96% ethanol and stirred for 1 h.

#### EC solution preparation

Various amounts of EC (10 g for 1:1 ratio, 20 g for 1:2 ratio, etc.) were dissolved in 400 ml of 96% ethanol, with 2.5% w/v propylene glycol added. Each solution was stirred with a magnetic stirrer for 3 h.

#### Mixing solutions

The amylose and EC solutions were mixed according to their specific ratios and stirred for an additional 6 h after adding erythrosine.

### Film coating of metronidazole core tablets

Metronidazole tablets were coated using a pan coater (ERWEKA Type: UG, Model: Nr.10736, ERWEKA-APPRATEBAU-GmbH, Heusenstamm Kr. Offenbach/Main, Germany) as described elsewhere [19, 20]. Coated samples with different film thicknesses (2%, 4%, and 6% total weight gain, TWG) were obtained by taking samples at intervals. Tablets were pre-warmed in the coating pan for 10 minutes at 40-45°C. Fifty tablets were randomly taken, weighed, and their average weight calculated for baseline TWG.

Coating solutions (1:0 to 1:5 ratios) were divided into three sub-batches to achieve 2%, 4%, and 6% coating thicknesses. After coating, the tablets were dried for an additional 10 minutes with hot air spray at 40-45°C. Coating parameters are shown in Table 2.

**Table 2.**
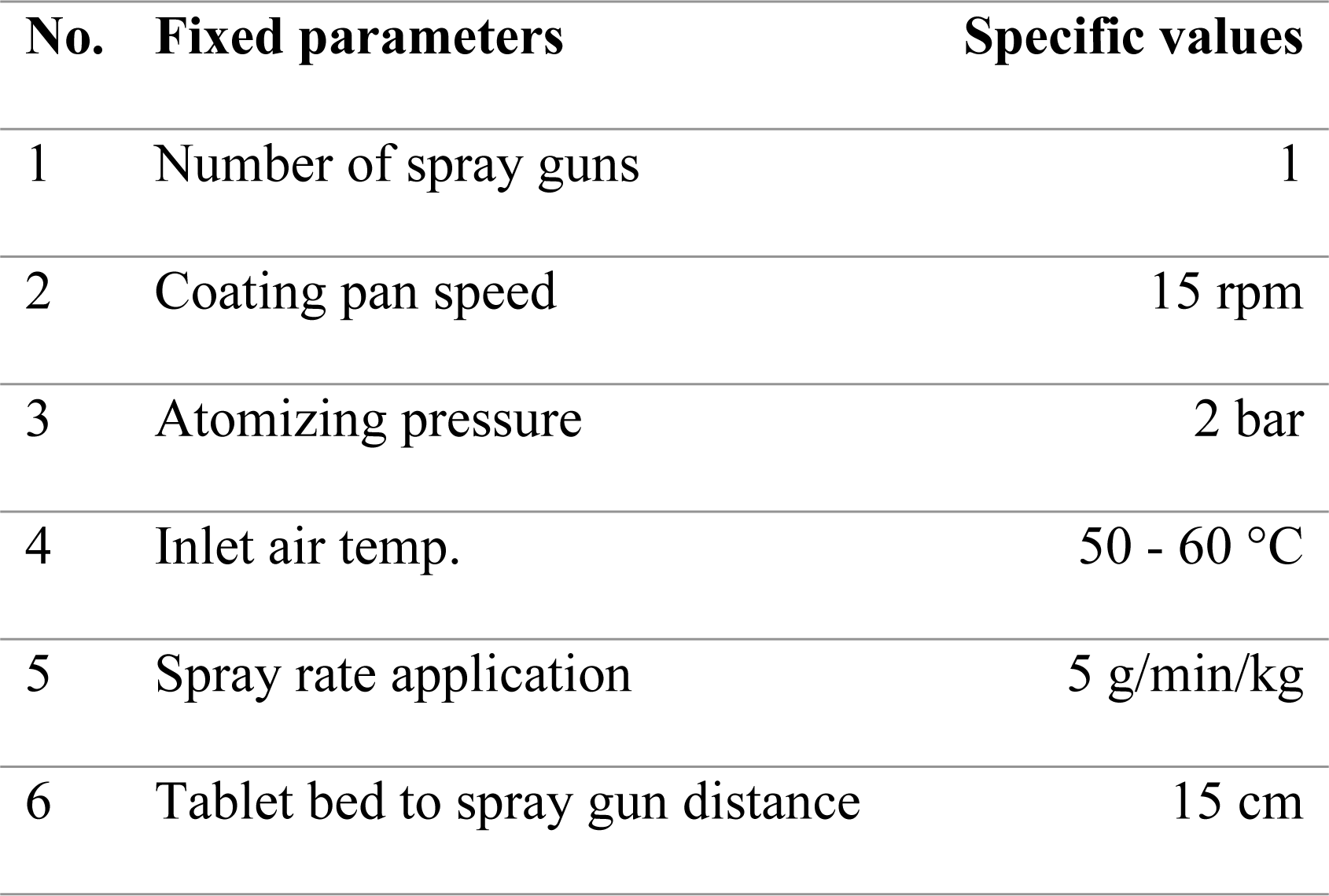
Coating process parameters that were maintained during the coating of tablets.

| No. | Fixed parameters | Specific values |
| --- | --- | --- |
| 1 | Number of spray guns | 1 |
| 2 | Coating pan speed | 15 rpm |
| 3 | Atomizing pressure | 2 bar |
| 4 | Inlet air temp. | 50 - 60 °C |
| 5 | Spray rate application | 5 g/min/kg |
| 6 | Tablet bed to spray gun distance | 15 cm |

### Evaluation of film coated tablets

#### Percentage weight gain of coated tablets

The thickness of the film coating was expressed as the percentage total weight gain (TWG), calculated using Eq. 1

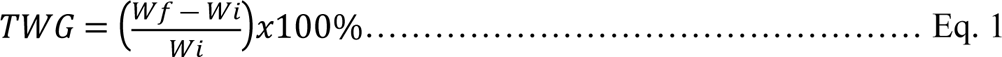

where *Wi* and *Wf* were initial and final tablet weights, respectively.

Three groups of film thicknesses (2%, 4%, and 6% TWG) were evaluated for each of the six different Amylose: EC ratios, resulting in a total of eighteen samples being prepared and assessed.

### Dissolution test

Metronidazole release from the film-coated tablets was assessed using an in vitro dissolution test with a USP type I basket dissolution apparatus (Model: LABINDIA DS 8000). The basket rotation speed was set at 100 rpm, and 900 ml of dissolution medium was used, which varied at different stages of the experiment [17].

For the first 3 h, the test was conducted in simulated gastric fluid (0.1 N HCl with 0.32% pepsin), followed by simulated small intestinal fluid (pH 6.8 phosphate buffer with 1% pancreatin) for the next 3 h (4, 18, 19). At predetermined time points, 5 ml samples were withdrawn from the gastric medium at 15 min, 30 min, 45 min, 1 h, 2 h, and 3 h, and from the small intestinal medium at 4 h, 5 h, and 6 h. Samples were filtered using Double Rings® 102 Medium filter paper (9 cm), suitably diluted, and analyzed by UV spectrophotometry at a λmax of 278 nm. For each Amylose to EC ratio (1:0 to 1:5) and each thickness (2%, 4%, and 6%), the experiments were performed in triplicate.

### Fermentation study

Metronidazole release from the film-coated tablets was evaluated under in vitro conditions simulating the human colon, both with and without colonic bacteria. This assessment was conducted from the 6th to the 14th hour. One tablet was introduced into each of four 100 ml batch culture fermenters inoculated with human feces (10% w/v). The fermenters were prepared by homogenizing freshly voided human feces from three healthy subjects in a phosphate-based buffer medium with pH 7.2 [21–23].

The fermenters were sealed in airtight jars previously made anaerobic using an anaerobic kit and aerobic condition indicators (Anaerobic container system with an indicator; BD GasPak™ EZ, Becton Dickinson and Company, 7 Loveton Circle, Sparks, MD 21152, USA). The jars were placed in a shaker-incubator (Stuart®, Orbital Incubator / SI500 / BIBBY SCIENTIFIC LIMITED, STONE, STAFFORDSHIRE, ST15 0SA, UK) at 37°C and shaken at 100 rpm. The anaerobic environment was maintained with the candle jar method (burning a candle in a closed jar to remove oxygen) and monitored with an anaerobic indicator that turned blue upon exposure to oxygen. A control experiment, using a buffer medium without feces, was run in parallel. Each experiment was conducted in triplicate. Two milliliters of samples were collected every 15 minutes for the first hour, then at hourly intervals for the next 8 hours. The samples were filtered through 0.2 µm filters prior to analysis for drug concentration by UV/Vis spectrophotometry at λmax of 278 nm [24]. The results were plotted to show cumulative percentage drug release versus time profiles for both gastric and small intestine dissolutions up to the 14th h.

### Data analysis

Statistical analysis was performed using Analysis of Variance (ANOVA) with the software Origin 6.0 (Origin Lab TM Corporation, USA). Tukey’s multiple comparison tests were used to compare individual differences. All data reported are averages of at least triplicate measurements, expressed as mean ± standard deviation (SD) with a 95% confidence interval. p-values < 0.05 were considered statistically significant.

## Results and Discussion

### Evaluation of the extracted TS and RS

#### Qualitative and quantitative tests of TS and RS

The yield and other properties of teff grains of different traits have been characterized elsewhere [15, 19]. The starch sample solutions from the four teff varieties (labelled D1 to D4) turned blue-black upon the addition of iodine solution, confirming the presence of starch. The yields of TS and RS obtained from different teff varieties are summarized in Table 3.

**Table 3.** Yields of TS and RS isolated from 100 g of teff grains.

| **Teff grain** | **Teff flour\*** | **TS** | **RS** |
| --- | --- | --- | --- |
|  | **% yield** | **% yield** | **% yield** |
| DZ-01-1681 | 81.28±1.4 | 37.20±1.2 | 18.07± 0.8 |
| DZ-01-99 | 79.04±1.3 | 39.10±1.1 | 19.54±1.1 |
| DZ-01-196 | 84.16±0.9 | 40.40±1.3 | 19.74±0.9 |
| DZ-CR-37 | 83.60±1.1 | 34.90±0.9 | 18.04±0.6 |
| *Mean % yield* | 82.02 | 37.90 | 19.10 |
| *SD of the mean* | 2.35 | 2.39 | 0.76 |

\*Sieved through 315 µm sieve, TS: Total starch, RS: Resistant starch

The results indicate that the mean yield of TS from different teff varieties is 37.90 ± 2.39%, while the mean yield of RS from TS is 19.10 ± 0.76% (Table 3). The RS yields for the four varieties were not significantly different, but the TS yields varied significantly among varieties (p < 0.05).

### Evaluation of granule properties of the formulated blends

For tablets to be uniform in weight and content, proper blending of the powder is essential to ensure consistent flow and compaction during the tableting process. The angle of repose and flow rate of the granules were measured at 27.46 ± 1.6° and 16.90 ± 1.15 g/sec, respectively. According to USP 30-NF25 <1174> (2007) (16), the granules exhibited excellent flow properties as the angle of repose fell within the acceptable range of 25°–30°.

The Carr’s Index (15) and Hausner Ratio (19) were found to be 13.08 ± 1.48% and 1.15 ± 0.02, respectively. These values indicate “GOOD” flow characteristics according to the generally accepted scale, with a CI range of 11–15% and an HR range of 1.12–1.18 [17].

### Physicochemaical properties of core-tablets

Table 4 presents the physicochemical properties of the core tablets, including diameter, thickness, weight variation, hardness, friability, disintegration time, and content uniformity.

**Table 4.** Physicochemical properties of Metronidazole core tablets.

| Batch Code | Diameter (mm) | Thickness (mm) | Weight-variation (mg) | Hardness (Kg/cm<sup>2</sup>) | Friability (%) | DT (min) N=6 | Content uniformity N=20 |
| --- | --- | --- | --- | --- | --- | --- | --- |
|  | N=10 | N=10 | N=20 | N=10 | N=20 |  |  |
| 01 | 11.74±0.02 | 4.47±0.09 | 503.64±2.85 | 128.80±5.18 | 0.36±0.01 | 2.19±0.23 | 107.10 |
| 02 | 11.71±0.06 | 4.46±0.03 | 503.63±3.50 | 128.10±8.30 | 0.31±0.02 | 1.92±0.19 | 106.27 |
| 03 | 11.73±0.01 | 4.40±0.04 | 504.41±3.53 | 129.50±5.19 | 0.40±0.01 | 2.30±0.43 | 108.08 |

The mean diameter and thickness of the tablets ranged from 11.71 ± 0.06 to 11.74 ± 0.01 mm and 4.40 ± 0.03 to 4.47 ± 0.09 mm, respectively. There were no significant differences in these parameters at the 0.05 level (p > 0.05). The tablet weights ranged from 503.63 ± 3.50 to 504.41 ± 3.53 mg. The average percent deviation for 20 tablets of each batch was less than 15% (less than ±30.0 mg) from the active ingredient (Metronidazole 200 mg), indicating compliance with dosage uniformity requirements according to USP 30 [17].

The hardness and friability of the core tablets ranged from 128.10 ± 8.30 to 129.50 ± 5.19 kg/cm² and 0.31% to 0.40%, respectively. These values demonstrate acceptable mechanical strength and mechanical stability, with friability less than 1%. The disintegration times for all three batches were less than 3 minutes, well below the 15-minute maximum disintegration time limit specified in the USP [17].

### Drug release characteristics of the film-coated tablets

#### Dissolution studies

The drug release profile studies were conducted in simulated gastric, intestinal, and colonic fluids to evaluate the film-coated tablets as colon-targeted drug delivery systems (CTDDS). While in vivo conditions for drug release are complex and challenging to fully replicate *in vitro* [24], employing various in vitro setups—such as exposing the dosage form to different chemical (pH), biological (enzymatic), and mechanical (agitation) stresses—provides valuable insights into the sensitivity of drug delivery systems under biological conditions [3]. This in vitro information helps approximate in vivo release characteristics.

The dissolution profile of the tablets is illustrated in Figure 1 as a three-dimensional (3D) graph, where the Z-coordinate represents the Amylose to EC ratio (1:0, 1:1, 1:2, 1:3, 1:4, and 1:5) for each % total weight gain (TWG) (2%, 4%, and 6%). The Y-coordinate shows the cumulative percentage of metronidazole release for each sample, and the X-coordinate indicates the time required for the total dissolution process.

**Figure 1.**
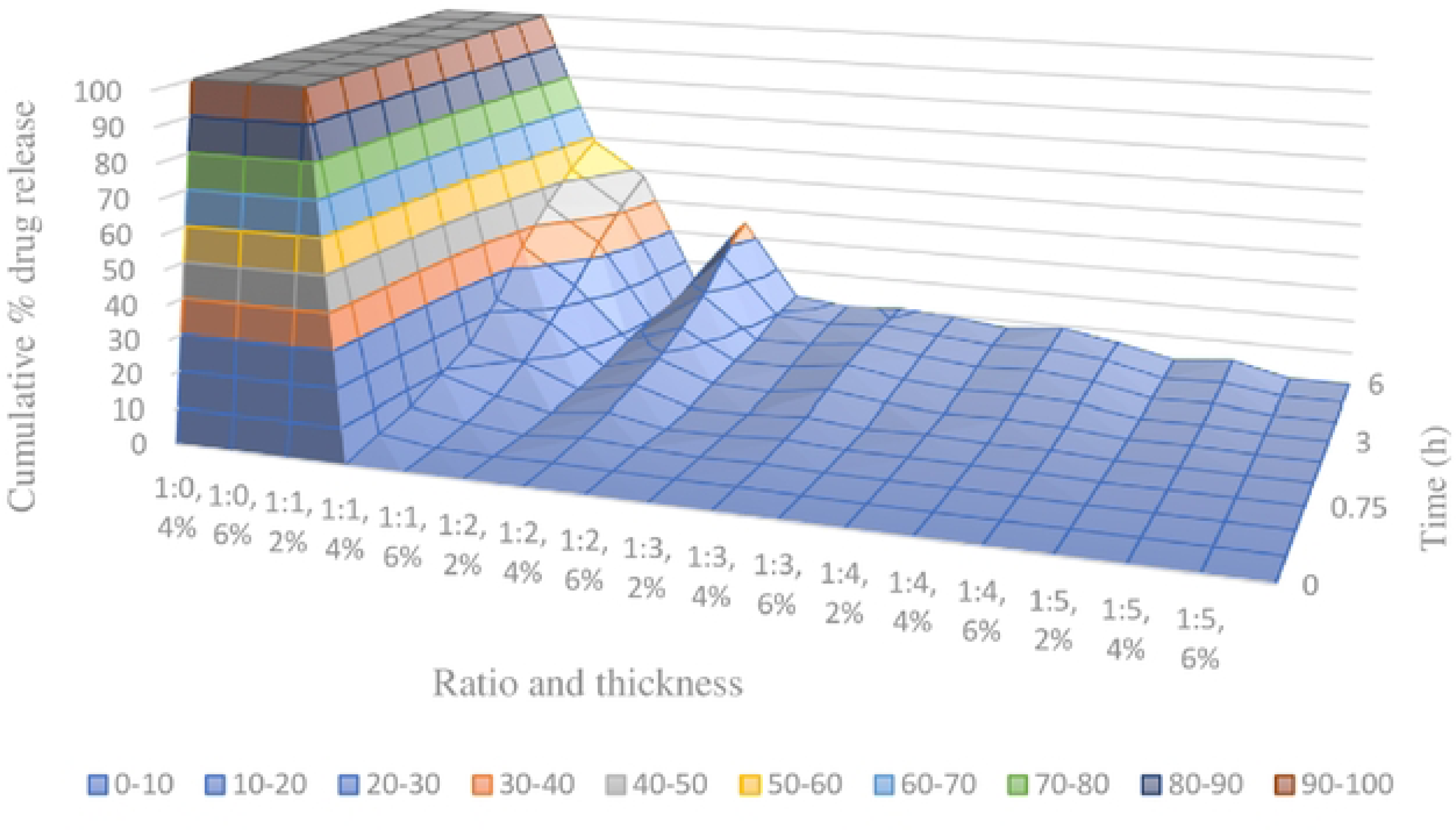
Dissolution profiles illustrating the cumulative percentage release of metronidazole from coated tablets with different amylose to EC ratios and coating thicknesses in simulated gastric fluid (from t = 0 to t = 3 h) and in simulated intestinal conditions (from t = 3 to t = 6 h).

A formulation for colon-targeted drug delivery should not release more than 15% of its content in the dissolution tests for the upper gastrointestinal tract (GIT)—i.e., in gastric and small intestinal fluids. This 15% cutoff is based on the USP 30 [17], which sp ecifies that not less than 85% of the labelled amount of metronidazole must be dissolved within 60 minutes for conventional tablet dosage forms. The CTDDS should ensure that at least 85% of the drug reaches the colon within the normal GIT transit time, implying that no more than 15% should be released in the upper GIT within 6 h.

Formulations that released more than 15% of their content in simulated gastric and small intestinal fluids were excluded as candidates for colon-targeted delivery. This includes all 2%, 4%, and 6% TWG thicknesses of the 1:0 amylose to EC ratio formulation, 2% and 4% TWG thicknesses of the 1:1 amylose to EC ratio, and 2% TWG thickness of the 1:2 amylose to EC ratio. The remaining 12 formulations were further evaluated in the fermentation test to assess their potential as colon-targeting systems.

The 1:0 polymer ratio formulation exhibited immediate release of metronidazole due to the readily swellable nature of amylose, which disrupted the film’s integrity and caused abrupt drug release. This issue was mitigated by incorporating the water-insoluble polymer EC to retard premature drug release. In other formulations (1:1 and 1:2 ratios of amylose to EC), the premature release in gastric and small intestinal media was attributed to their thin coating thicknesses and higher proportion of amylose in the film-forming material.

### Fermentation studies

Metronidazole release from the film-coated tablets was assessed under colon-simulating conditions (from the 6th to the 14th hour) at a pH around 7.2, with and without bifidobacteria cultured from the fecal matter of three healthy individuals. The results are presented as cumulative percentage drug release versus time profiles in Figures 2 and 3.

**Figure 2.**
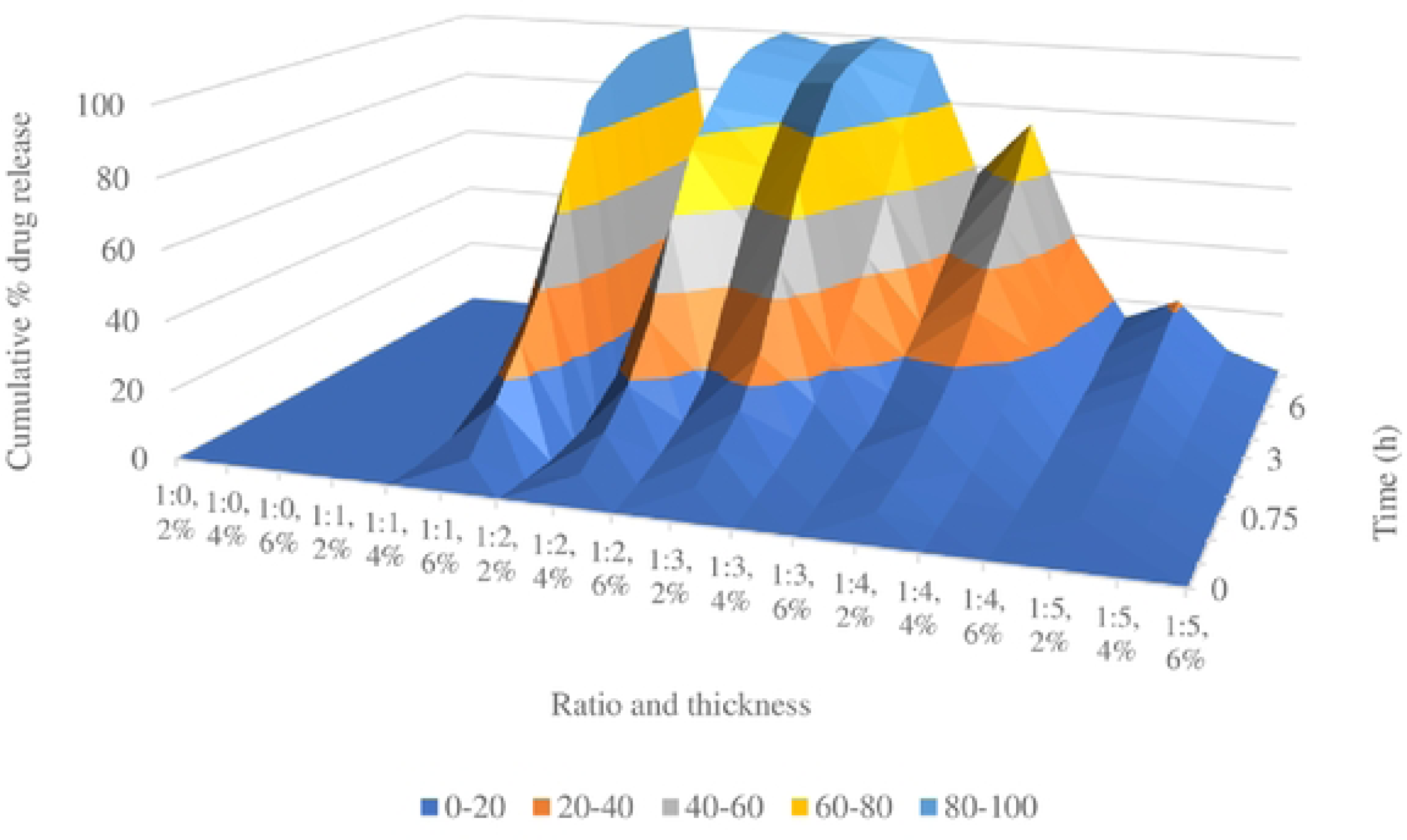
Metronidazole release from the film-coated tablets was assessed under colon-simulating conditions (from the 6th to the 14th hour) at a pH around 7.2, with and without bifidobacteria cultured from the fecal matter of three healthy individuals. T

**Figure 3.**
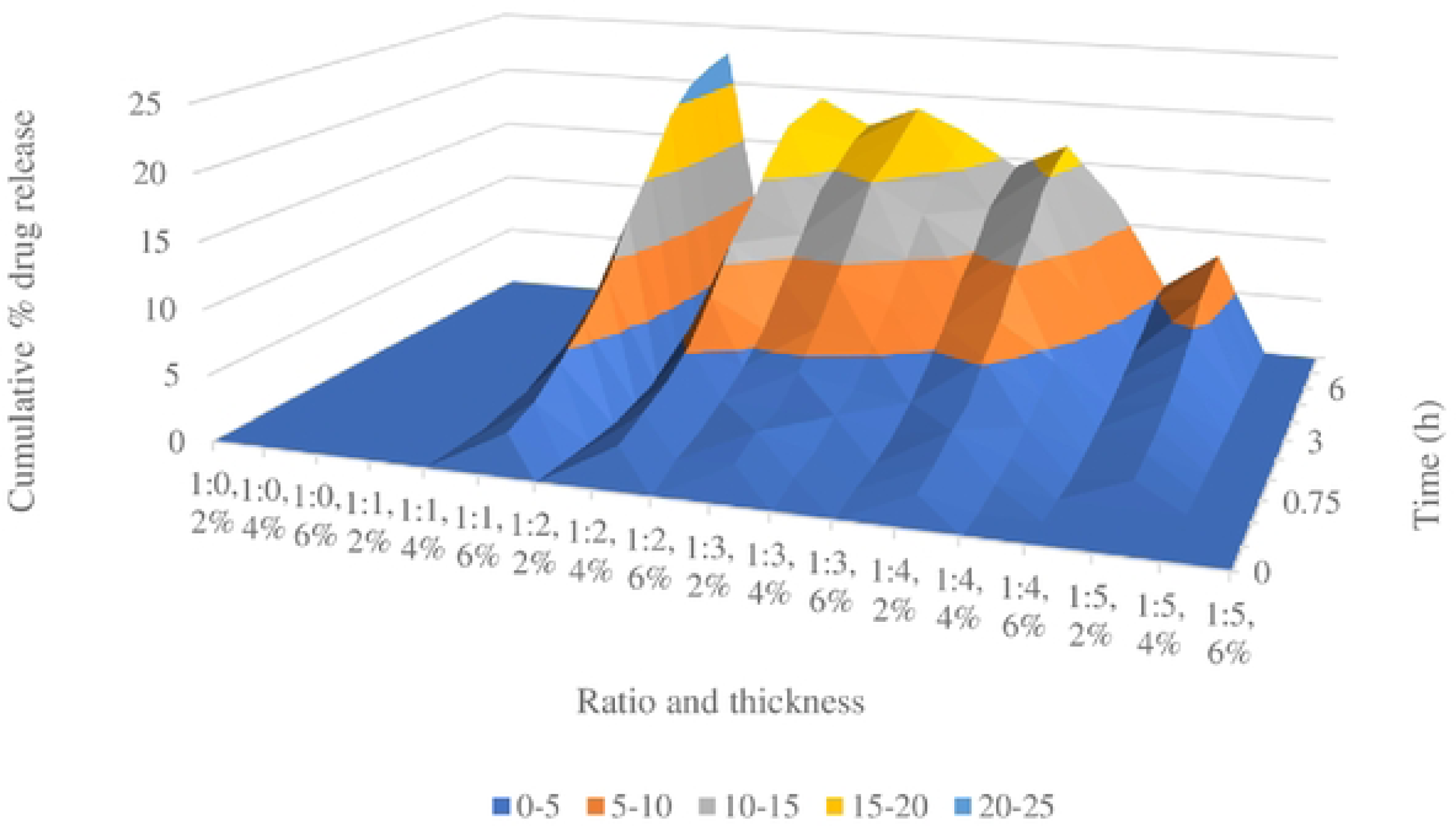
Release of metronidazole from film-coated tablets in colon-simulated fermentation medium without colonic bacteria, highlighting the influence of film-forming material ratios and coating thickness on drug release in a 3D graph.

Figure 2 provides a 3D graph for the remaining 12 samples, showing the results of the fermentation process over time (X-axis), cumulative percentage drug release (Y-axis), and the amylose to EC ratios (from 1:0 to 1:5) with coating thicknesses (2%, 4%, and 6%) (Z-axis).

According to the USP30 [17], a formulation is considered acceptable if it releases 85 ± 5% of the drug content. In Figure 2, five formulations that released more than 90% of the drug were deemed suitable CTDDS candidates: 1:1 ratio with 6% TWG, 1:2 ratios with 4% and 6% TWG, and 1:3 ratios with 2% and 4% TWG. In these formulations, drug content was released primarily in the colon-simulated medium, as the tablets either passed through the upper GIT intact or released only a small amount (≤ 10%) of their drug content. This localized release is due to the presence of colonic bacteria, which degrade the RS in the film coat, creating openings that facilitate drug release in the colon.

The drug release was found to decrease as the proportion of EC increased in the film, due to EC’s role as a barrier to drug release. EC is not digested by colonic bacteria and does not dissolve in the medium. Additionally, the thickness of the film coat inversely affected drug release; thicker films delayed drug exposure as they required more time for bacterial digestion of RS and had longer diffusion paths for drug release.

To confirm the role of colonic bacteria in drug release, the fermentation experiment was repeated without human feces (i.e., in the absence of colonic bacteria). The results, shown in Figure 3, display cumulative percentage release (Y-axis) for each sample thickness in % TWG (2%, 4%, and 6%) with varying amylose to EC ratios (Z-axis) over time (X-axis).

In Figure 3, all tested formulations released no more than 25% of their content, regardless of film thickness and amylose to EC ratio. This result underscores the critical role of colonic bacteria in drug release from the CTDDS. Without fecal matter, the absence of bacteria meant that RS was not digested, preventing pore formation and drug release. Thus, the amount of amylose and bacterial substrate are crucial factors impacting drug release, while film thickness primarily influences the diffusion path length.

### Drug release: summary of dissolution and fermentation studies

The drug release from film-coated tablets is influenced by both diffusion and erosion mechanisms. In simulated gastric and intestinal environments, drug release is primarily controlled by diffusion through pores, as film cracks occur before significant erosion. The larger volume of fluid in the upper GIT facilitates this diffusion.

In the colonic environment, drug release is governed by the superposition of diffusion and erosion. Due to the smaller volume of colonic fluid, diffusion becomes less significant. Instead, once the film coat, composed of natural polysaccharides, is fermented by colonic microflora, the resulting erosion predominates [5, 25, 26].

Factors such as GI motility, disease conditions, and colonic bacterial count may influence drug release from bacterially triggered CTDDS [3]. Assuming these factors are constant or vary minimally, the results of the dissolution and fermentation studies indicate that the ratio of amylose to ethylcellulose (EC) in the film coat and the coating thickness are critical parameters for the CTDDS developed. The optimum drug release percentages were found between the two extreme values tested, with formulations exhibiting the best balance located around the center of these extremes (Table 5).

**Table 5.**
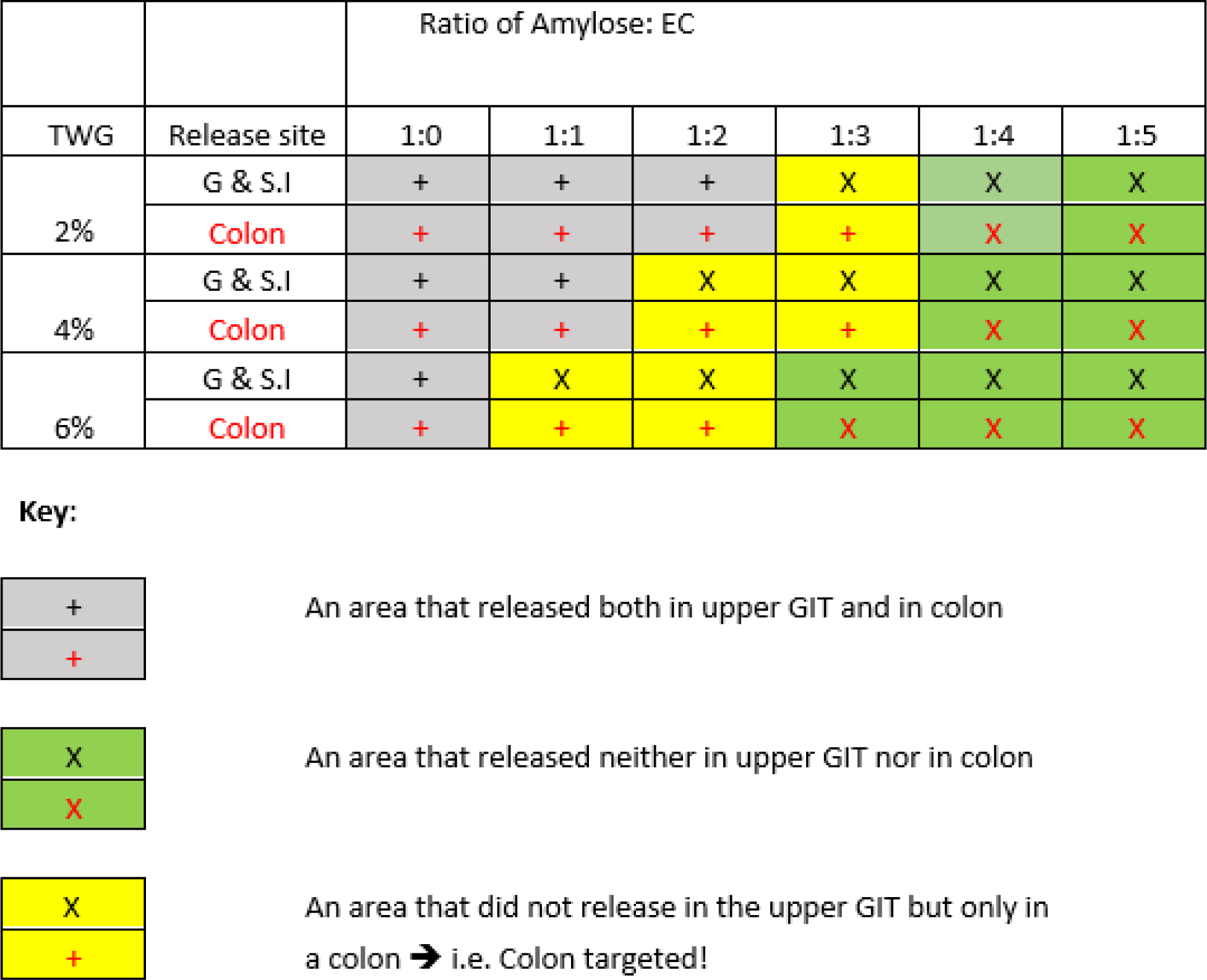
Summary of dissolution and fermentation studies for different ratio of amylose to EC with six sample groups each having thickness of 2%, 4% and 6% in simulated gastric, small intestinal and colonic environments.

## Conclusion

The study isolated starch and resistant starch (RS) from local teff grains and utilized the latter to develop a film-forming coating for colon-targeted drug delivery systems (CTDDS). Through a series of in vitro dissolution and fermentation studies, it was determined that the ratio of amylose to ethylcellulose (EC) in the film coating material played a crucial role in controlling the drug release process, while the thickness of the film was a secondary factor. The optimal combinations identified for effective drug release in the colon, without premature release in the upper gastrointestinal tract (GIT) were: 1:1 amylose to EC ratio with 6% film thickness; 1:2 amylose to EC ratio with 4% and 6% film thickness; and 1:3 amylose to EC ratio with 2% and 4% film thickness. These formulations demonstrated the ability to restrict drug release to the colonic environment while minimizing release in the upper GIT. The results highlight the potential of using RS isolated from local teff grains as a viable excipient for CTDDS, offering an effective and localized drug delivery approach. In summary, the study underscores the effectiveness of incorporating RS from teff grains into film coating materials for CTDDS and identifies the key parameters for optimizing drug release profiles.

## Acknowledgements

The authors would like to thank the Ethiopian Pharmaceuticals Sh. Co. for providing access to their lab facility, and for the provision of API, and chemicals, the Armauer Hansen Research Institute for access to their lab facility and donation of materials for the fermentation process, and the Holeta Agricultural Research Institute for providing the four varieties of teff grains. YT acknowledges Addis Ababa University for sponsoring his MSc research.

